# Proton-Dependent Inhibition, Inverted Voltage Activation, and Slow Gating of CLC-0 Chloride Channel

**DOI:** 10.1101/2020.10.02.323352

**Authors:** Hwoi Chan Kwon, Yawei Yu, Robert H. Fairclough, Tsung-Yu Chen

**Author notes:** Department of Molecular Biology, University of California, Berkeley, California, United States of America (Yawei Yu). Corresponding Author: (TC).

## Abstract

CLC-0, a prototype Cl^−^ channel in the CLC family, employs two gating mechanisms that control its ion-permeation pore: fast gating and slow gating. The negatively-charged sidechain of a pore glutamate residue, E166, is known to be the fast gate, and the swinging of this sidechain opens or closes the pore of CLC-0 on the millisecond time scale. The other gating mechanism, slow gating, operates with much slower kinetics in the range of seconds to tens or even hundreds of seconds, and it is thought to involve still-unknown conformational rearrangements. Here, we find that low intracellular pH (pH_i_) facilitates the closure of the CLC-0’s slow gate, thus generating current inhibition. The rate of low pH_i_-induced current inhibition increases with intracellular H^+^ concentration ([H^+^]_i_)—the time constants of current inhibition by low pH_i_ = 4.5, 5.5 and 6 are roughly 0.1, 1 and 10 sec, respectively, at room temperature. In comparison, the time constant of the slow gate closure at pH_i_ = 7.4 at room temperature is hundreds of seconds. The inhibition by low pH_i_ is significantly less prominent in mutants favoring the slow-gate open state (such as C212S and Y512A), further supporting the fact that intracellular H^+^ enhances the slow-gate closure in CLC-0. A fast inhibition by low pH_i_ causes an apparent inverted voltage-dependent activation in the wild-type CLC-0, a behavior similar to those in some channel mutants such as V490W in which only membrane hyperpolarization can open the channel. Interestingly, when V490W mutation is constructed in the background of C212S or Y512A mutation, the inverted voltage-dependent activation disappears. We propose that the slow kinetics of CLC-0’s slow-gate closure may be due to low [H^+^]_i_ rather than due to the proposed large conformational change of the channel protein. Our results also suggest that the inverted voltage-dependent opening observed in some mutant channels may result from fast closure of the slow gate by the mutations.

## Introduction

The CLC channel/transporter family consists of transmembrane proteins of two functional categories: Cl^−^ channels and Cl^−^/H^+^ antiporters [1, 2]. These CLC proteins are expressed in various tissues to carry out critical physiological functions [3]. CLC-0, for example, is a Cl^−^ channel expressed in *Torpedo* electroplax [4], and the *Torpedo* fish exploits CLC-0’s function in the electric organ for building an under-water stun gun. CLC-1, CLC-2, and CLC-Ks are mammalian CLC channels important for normal functions of various organs, such as skeletal muscles, kidney, heart and brain [5–7]. On the other hand, several bacterial CLC molecules, such as CLC-ec1 [8], function as Cl^−^/H^+^ antiporters [9], which enable bacteria to develop resistance to H^+^ entry to the cells in very acidic environments [10]. Mammalian CLCs other than those mentioned above, such as CLC-5 [11], also function as Cl^−^/H^+^ antiporters [12]. They are thought to be important in controlling the pH in the intracellular organelles, and mutations of these CLC proteins have been known to be associated with human hereditary diseases such as Dent’s disease, osteomalacia, and lysosomal storage diseases [13–15].

It is known that all CLC members are homodimers [8, 16], and recent efforts have unveiled their molecular structures [17–22]. An example of the structure of a CLC protein most homologous to CLC-0 (i.e., CLC-1) is shown in Fig. 1 (PDB accessing code: 6QVU). Because of its well-characterized functional behaviors, CLC-0 is viewed as a prototype CLC molecule among CLC family members. Early single-channel recordings suggested the presence of two identical Cl^−^-conducting pores in CLC-0 [23–25], a double-barreled architecture later confirmed by CLC proteins’ high-resolution structures. Two gating mechanisms have been identified in controlling the opening and closing of CLC-0: “fast gating” and “slow gating.” Fast gating controls the two pores independently, and operates on the millisecond time scale, while slow gating operates on the order of ~seconds to hundreds of seconds [26]. Because the slow-gating mechanism appears to control the two pores simultaneously, it is also called “common” gating, and the closure of the slow gate “inactivates” the channel. Based on single-channel behaviors of CLC-0, when the slow gate closes, Cl^−^ conduction through the channel pores is shut, and the functional activities of the fast gate are not observable.

**FIGURE 1:**
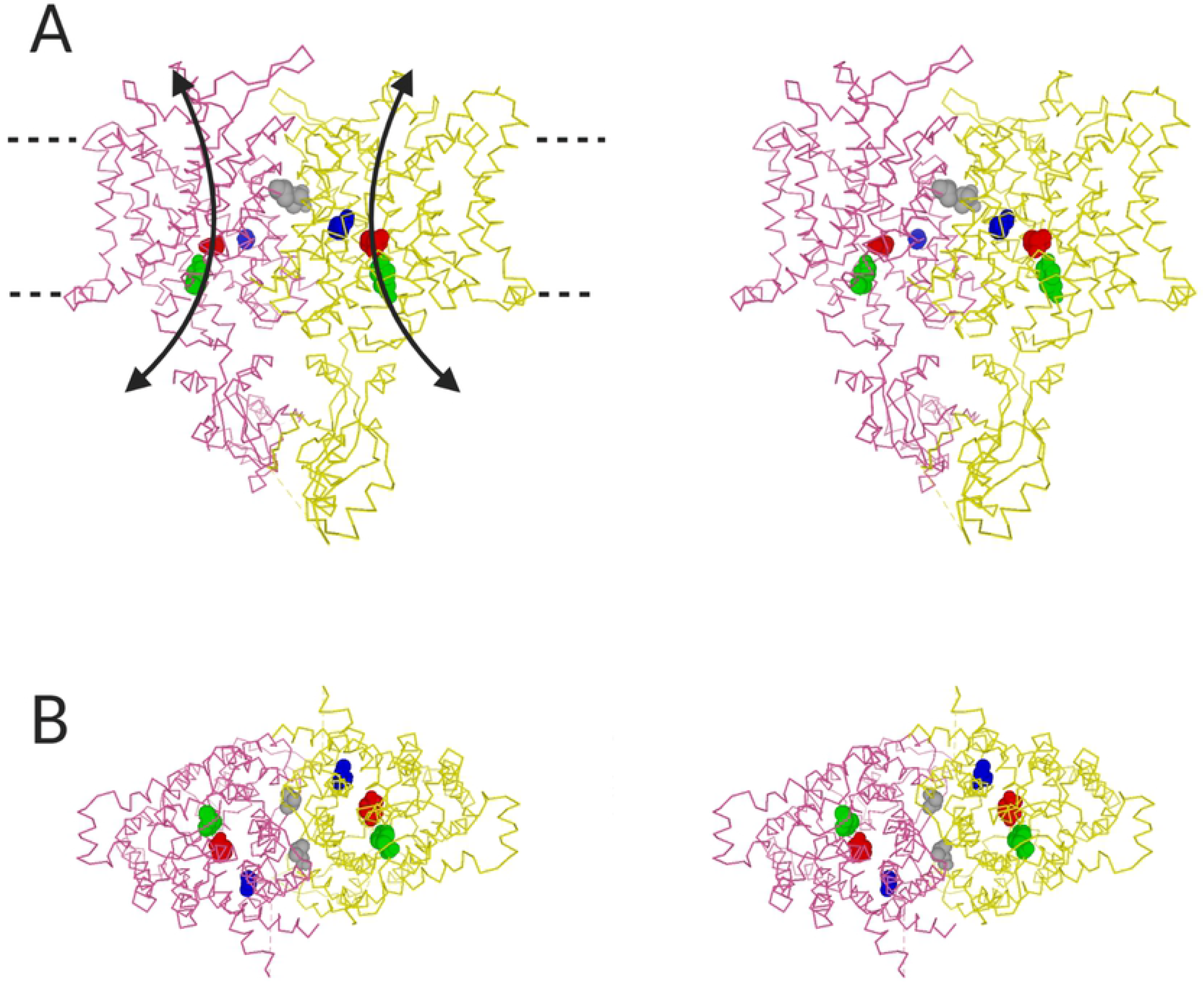
Structure of vertebrate CLC channels. Structure of human CLC-1 molecule (PDB accessing code: 6QVU) is used to represent the structure of CLC-0, which is still not available. CLC-0 residues mutated in this study are depicted by the colored and space-filled corresponding CLC-1 residues (in parenthesis): Blue, C212 (C277 of CLC-1); Grey, V490 (I556); Green, Y512 (Y578). The “E-gate” residue, E166 of CLC-0 (E232 of CLC-1), is also shown as a space-filled residue (red). **(A)** Stereo-view of hCLC-1 structure viewed from within membrane phospholipids (side view). Curved arrows depict the ion permeation pathways. Dotted lines indicate the extracellular and intracellular edges of lipid membranes. **(B)** Stereo-view of hCLC-1 viewed from the cytosolic side.

Various studies in the literature have characterized CLC-0’s fast-gating details. It has been shown that Cl^−^ in the extracellular and intracellular solutions both favor the opening of the fast gate of CLC-0 [27–29]. High-resolution CLC structures reveal three Cl^−^-binding sites in the ion-transport pathways of CLC proteins: the external (S_ext_), central (S_cen_) and internal sites (S_int_) [18]. However, S_ext_ can also be occupied by the negatively charged sidechain of a glutamate residue (corresponding to E166 in CLC-0), which is thought to be the fast gate (called E gate). Swinging this E-gate away from S_ext_ is considered to be the fast-gate opening mechanism, and thus facilitating the fast-gate opening by Cl^−^ could be due to a competition of Cl^−^ with the E-gate [30–32]. Extracellular and intracellular low pH also favors fast-gate opening [33–36], presumably due to the protonation of the E-gate.

Compared to fast gating, the molecular mechanism of CLC-0’s slow-gating is less defined despite being functionally characterized. Unlike the fast-gate opening, which is favored by membrane depolarization [24, 25, 27, 28], the voltage-dependence of slow-gate opening is opposite—the slow-gate’s open probability (P_o_^s^) decreases with membrane depolarization but increases with membrane hyperpolarization [37–39]. Cl^−^ and H^+^ also modulate CLC-0’s slow gating [40, 41] and the slow gating is found to be very temperature-dependent [38, 39]. Because mutations at multiple sites on the channel protein alter the slow-gating [42–45], it has been thought that slow gating may involve a large protein conformational change, including relative movement of the two subunits at both the cytoplasmic and the transmembrane regions [43, 46].

Because defective functions of mammalian CLC channels underlie human diseases, understanding CLC channel’s gating mechanisms is clinically relevant. For example, CLC-1 channelopathy causes a hereditary muscle disease, myotonia congenita. The myotonia pathophysiology is rationalized from the fact that CLC-1 constitutes 50-70 % of the resting muscle conductance and thus is critical for controlling sarcolemmal potential [47]. Similar to CLC-0, membrane depolarization favors the opening of CLC-1. A defect in the CLC-1 opening by depolarizing voltage therefore renders it difficult to bring the membrane potential back to the resting level after firing action potentials [48], thus generating a myotonia condition. Indeed, some CLC-1 myotonia mutants are opened by membrane hyperpolarization but not by depolarization [49, 50]. Such an inverted voltage dependence of channel opening also occurs in mutants of CLC-0. Using a voltage protocol shown in Fig. 2 A, for example, the voltage-dependent opening of wild-type (WT) CLC-0 and that of a point mutant, V490W, are compared in Fig. 2 B, where the current of the V490W mutant is activated by membrane hyperpolarization but not by depolarization. Interestingly, we discover that such a hyperpolarization-induced channel opening can also occur in WT CLC-0 in the presence of low pH_i_ (Fig. 2 C). In this paper, we study the relation between the inverted voltage-dependent channel opening and the intracellular H^+^ effect on the slow gating in CLC-0.

**FIGURE 2:**
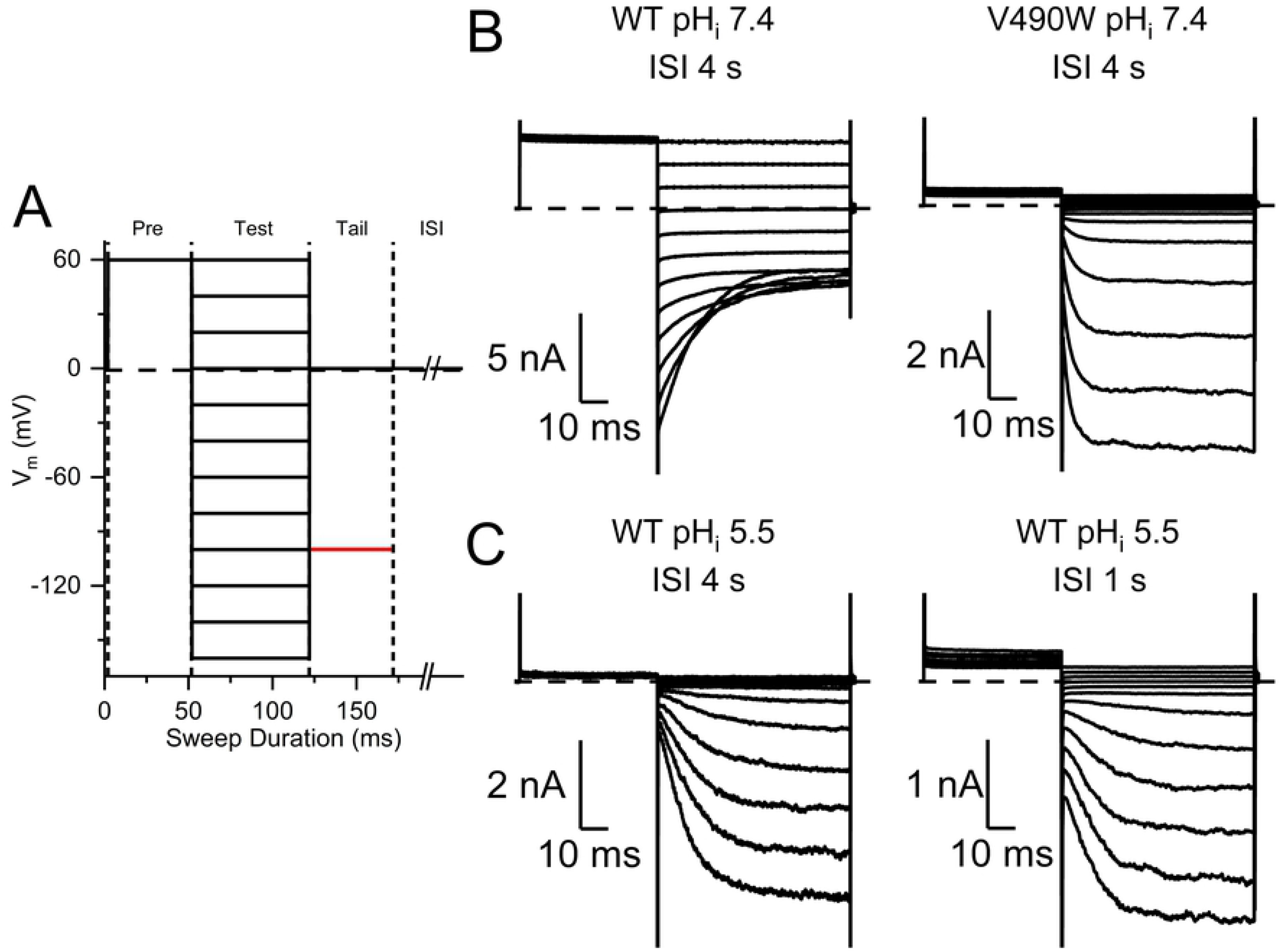
Voltage dependence of current activation of WT and mutant CLC-0. **(A)** Voltage protocol (protocol I) for recordings. A full protocol consists of 12 recording sweeps. One sweep includes a prepulse voltage step at +60 mV (50 ms) followed by one of the various test voltage steps (70 ms) from +60 mV to −160 mV in −20 mV voltage steps, and followed by a tail voltage step of 50 ms at 0 mV (colored in black) or −100 mV (colored in red). The voltage of the inter-sweep interval (ISI) was 0 mV and the duration was 1 or 4 sec. **(B)** Activation of WT CLC-0 and V490W mutant at pH_i_ = 7.4, using the voltage protocol shown in A. ISI was 4 sec. Dash line: zero-current level. Notice the inverted voltage-dependent activation in the mutant. **(C)** Activation of WT CLC-0 at pH_i_ = 5.5. ISI’s are 4 and 1 sec for the recording on the left and right, respectively. Notice the inverted voltage activation of WT CLC-0 in low pH_i_.

## Materials and methods

### Mutagenesis and channel expression

The cDNAs of the WT CLC-0 and various mutants of CLC-0 were subcloned in the pIRES2-EGFP vector containing internal ribosome entry sites (IRES) and enhanced green fluorescent protein (EGFP). Mutagenesis was made using the Quick Change site-directed mutagenesis kit (Strategene), and the mutations were confirmed via commercially available sequencing services. All cDNAs were transfected into the human embryonic kidney 293 (HEK293) cells grown in Dulbecco’s modified Eagle’s medium (DMEM) supplemented with 10% fetal bovine serum (FBS) and 1% Penicillin/streptomycin. The transfections were performed using commercially available Lipofectamine 3000 kit (Invitrogen) following the standard protocol provided by the vendor. After transfections, cells were incubated in 37 °C and 5% CO_2_ for 1-2 days before conducting experiments.

### Electrophysiological Recordings

Transfected HEK293 cells were identified by the green fluorescence under the Leitz DM IRB inverted microscope (Leica) equipped with GFP filter (Chroma Technology) and the XT640-W LED light source (Lumen Dynamics). Inside-out membrane patches were excised from the green fluorescence-positive cells, and voltage-clamp experiments were conducted using the Axopatch 200B amplifier (Axon Instruments/Molecular Devices). The recorded signals were filtered at 2 kHz and were digitized at 4 kHz using Digidata 1440A digitizing board (Molecular Devices/Axon Instruments). Occasionally, the 50/60 Hz noise signal was removed using Hum Bug 50/60 Hz eliminator (Quest Scientific). Recording pipettes were fabricated from the TW150-6 borosilicate glass capillaries (World Precision Instruments Inc.) using the pp830 vertical puller (Narishige International). The electrode resistance was normally ~ 2-3 MΩ when filled with the pipette (extracellular) solution containing 130 mM NaCl, 5 mM MgCl_2_, 10 mM HEPES, 1 mM EGTA, with the pH adjusted to 7.4. The intracellular solution is the same as the extracellular solution except that 10 mM MES was used as the pH buffer when pH < = 6.2.

Fast solution exchange was achieved using the SF-77B solution exchanger (Warner Instruments/Harvard apparatus). Although the time for crossing the laminar flow barrier is estimated to be several ms (Zhang et al., 2009), there is always a latency time of 20-50 ms due to the time lag in initiating the crossing. Therefore, time constants of less than 50 ms estimated from a current relaxation following a change of pH_i_ were considered less accurate. Most of the experiments were performed at room temperature (21-22 °C). When a higher temperature was required, the solutions in the 10 ml syringe reservoirs were raised to a constant higher temperature using the SW-10/6 multi-syringe warmer alongside with the TC-324B controller (Warner instruments/Harvard Apparatus). The temperature of the solution, however, dropped when it was flowing through the PE50 tubing and finally exited out of the SF-77 barrel tip where the solution meets the excised inside-out patch. The reported temperatures in this study were those recorded by a thermistor placed at the outlet of the SF-77 solution delivery barrel after every recording experiment. It should be emphasized that such a temperature control was not optimal, and the temperature variation can be up to 1-2 ^o^C based on the variation among multiple measurements with the same solution temperature in the syringe reservoir.

### Experimental protocols and Data Analyses

Because the slow-gate opening of CLC-0 tends to be minimal at voltages of the resting membrane potential of HEK293 cells or above (Chen 1998), WT CLC-0 was partially inactivated when membrane patches were excised. To activate the current of CLC-0 for experiments, five 50-ms pulses of −100 mV at 1 Hz were applied to all patches expressing WT CLC-0 before starting any experiment. The procedure was repeated multiple times until the slow gate was maximally opened (based on the observation that the recorded current was no longer increased by this current-activation procedure). For mutants with a mostly open slow-gate (such as C212S or Y512A) no such current activation procedure was necessary.

Three types of experimental protocols were employed for the experiments presented in this paper. Protocol I was used for evaluating the quasi steady-state voltage-dependent current activation. Each recording sweep in this protocol has a total time of 170 ms, consisting of a +60 mV pre-pulse voltage step for 50 ms, followed by a test voltage step (from +60 mV to –160 mV in a series of −20 mV voltage steps) for 70 ms, and finally a tail voltage (at 0 mV or −100 mV) for 50 ms. The inter-sweep interval (ISI), namely, the time between the end of one recording sweep to the beginning of next sweep was 1 sec or 4 sec. Membrane voltage of ISI was at 0 mV. When this experimental protocol with a - 100 mV tail-voltage was employed, analyzing the initial tail current (obtained at the beginning of the - 100 mV tail voltage step) provides an estimate of the relative open probability (P_o_) of the channel [43]. In WT CLC-0 recordings, the largest initial tail current (negative current) occurred in the recording sweep with the +60 mV test voltage step because the overall channel opening is favored by membrane depolarization. On the other hand, for the mutant activated by membrane hyperpolarization (such as V490W), the largest tail current was observed in the recording sweep with the −160 mV test voltage [43]. In either case, the tail-current relaxation process was fitted to a single-exponential function, and the initial tail current from each recording sweep was determined by extrapolating the exponential tail current relaxation to the beginning of the tail voltage step. The initial tail currents from all recording sweeps were normalized to the maximal initial tail current, which represents a relative P_o_ of the channel at the end of the test voltages (relative to the P_o_ at the most positive voltage in the WT CLC-0 or to the most negative voltage in the mutant V490W). Plotting the relative P_o_ as a function of the test voltages illustrates the voltage dependence of the channel activation.

To evaluate the process of the change of Cl^−^ current upon reducing pH_i_, we employed a voltage protocol (protocol II) containing a voltage step of +60 mV (50 ms) followed by a tail voltage step at - 100 mV for 70 ms. Such a voltage protocol was used to mimic the experimental protocol of the previous studies using whole cell recording methods on channels expressed in Xenopus oocytes [38, 39]. To present the experimental results, the current at the +60 mV voltage step was measured and plotted against the time of the recording. ISI, which was either 1 or 4 sec, also refers to the time interval between the end of one recording sweep and the beginning of the following sweep. The membrane voltage at the ISI in this protocol was also 0 mV.

The third voltage protocol (protocol III), like protocol II, was also used to assess the current inhibition process after applying a high [H^+^]_i_ except that the application and removal of the low pH_i_ solution was conducted at a constant membrane voltage. For these experiments, the membrane voltages ranged from +60 mV to −60 mV. However, experiments with a slow inhibition process (in relatively high pH_i_ conditions such as pH_i_ = 5.5 or 6) at some negative voltages were technically difficult due to stability problems of the excised patches. Estimates of the time constant of such slow inhibition processes may thus be less precise.

Protocol II & III were used for studying the kinetics of the current relaxation process followed by the pH_i_ perturbation. Both protocols started from a steady-state current level at pH_i_ = 7.4. A lower pH_i_ solution was then applied, and the current was inhibited to different degrees at different speeds depending on pH_i_ (or [H^+^]_i_). After the current reached a steady state, pH_i_ was changed back to 7.4 and the current may or may not recover. The current relaxation process (current inhibition or current recovery) was then fitted to a single-exponential function to obtain the time constant of current inhibition (τ_inh_) and the time constant of current recovery (τ_rec_). However, for recordings of the WT CLC-0, current recovery was observed only at negative membrane voltages but not at the positive membrane voltages.

## Results

The fast and the slow gating of CLC-0 have opposite voltage dependence. Membrane depolarization increases the fast-gate open probability (P_o_^f^) but reduces the slow-gate P_o_(P_o_^s^). Using protocol I (Fig. 2 A) in which the duration of a recording sweep is less than 0.2 sec, the change of P_o_^s^ of CLC-0, due to its very slow kinetics (tens to hundreds of seconds at room temperature), does not result in a dramatic current change in such a short time frame. Therefore, the change of the WT CLC-0 current shown in Fig. 2 B (left panel) mostly reflects the activities of the fast gating. When the membrane voltage is at 0 mV or above, P_o_^f^ of CLC-0 is maximal (P_o_^f^ ~ 1) [25, 28, 44, 51]. Therefore, the instantaneous current jump upon a change of the voltage is due to the change of the driving force rather than an alteration of channel’s P_o_^f^. On the other hand, when membrane voltage is hyperpolarized, P_o_^f^ is reduced, and a current reduction with a deactivation time constant of several milliseconds is observed (Fig. 2 B, left panel). The current deactivation at hyperpolarization voltages reflects a reduction of P_o_^f^, with a kinetics in the millisecond time range.

In some mutant channels of CLC-0, such a normal voltage dependence of channel opening is inverted. For example, using the same voltage protocol I (Fig. 2 A), WT CLC-0 (Fig. 2 B, left panel) and the V490W mutant (Fig. 2 B, right panel), have opposite voltage dependence for current activation [43]. In the V490W mutant, little current is observed at voltages > 0 mV, while membrane hyperpolarization activates the current. Thus, membrane hyperpolarization but not depolarization favors the opening of this mutant channel. Interestingly, when pH_i_ is reduced, even a WT CLC-0 channel can exhibit such an inverted voltage-dependent channel activation. Fig. 2 C shows that at pH_i_ = 5.5, voltage steps above 0 mV activate little current in WT CLC-0, while membrane hyperpolarization induces current similar to that observed in the V490W mutant at a neutral pH_i_.

When protocol I was used to activate the current of WT CLC-0 at low pH_i_, the duration of the inter-sweep interval (ISI) (where the membrane voltage was held at 0 mV) played an important role for the amplitude of the activated current. This can be observed by comparing the two recordings of WT CLC-0 at pH_i_ = 5.5 obtained with the same voltage protocol (protocol I) but different ISI durations: 4 sec *versus* 1 sec (Fig. 2 C left and the right panels, respectively). Comparing the current at the +60 mV pre-pulse voltage step, the recording with 4-sec ISI (left panel) shows little outward current while the pre-pulse current in the recording with 1-sec ISI retains some outward current. The difference in the outward current between these recordings is also reflected by the instantaneous inward current when the membrane voltage is changed from +60 mV pre-pulse voltage step to the various hyperpolarizing voltage steps. It should be noted that the current at the +60 mV pre-pulse voltage depends on the channel conductance at the end of the ISI following the previous recording sweep. We suspected that intracellular H^+^ may inhibit the current of WT CLC-0 at the 0-mV holding voltage during ISI. Therefore, in a recording with 4-sec ISI, the channel conductance (after being activated by membrane hyperpolarization) was nearly completely inhibited by H^+^. On the other hand, with 1-sec ISI, the hyperpolarization-activated conductance has not been completely inhibited before starting the following recording sweep.

To confirm this speculation, we employed a continuous recording protocol, in which a sweep of recording contains only a +60 mV voltage step for 50 ms followed by a negative tail voltage step of −100 mV for 70 ms (protocol II, Fig. 3 A). The holding voltage at ISI was 0 mV. Typical experiments are shown in Fig. 3 B & C, where the ISI is 4 and 1 sec, respectively. These experiments started with a steady-state recording at pH_i_= 7.4. An acidic intracellular solution (pH_i_ = 5 in these two experiments) was then applied, followed by a switch of solutions back to pH_i_ = 7.4. Each circle in Figs. 3 B & C represents the outward current measured at the end of +60 mV voltage step. These experiments show that the outward current in the recording with ISI = 4 sec (Fig. 3 B) is inhibited almost completely while the recording with ISI = 1 sec (Fig. 3 C) still retains significant outward current.

**FIGURE 3:**
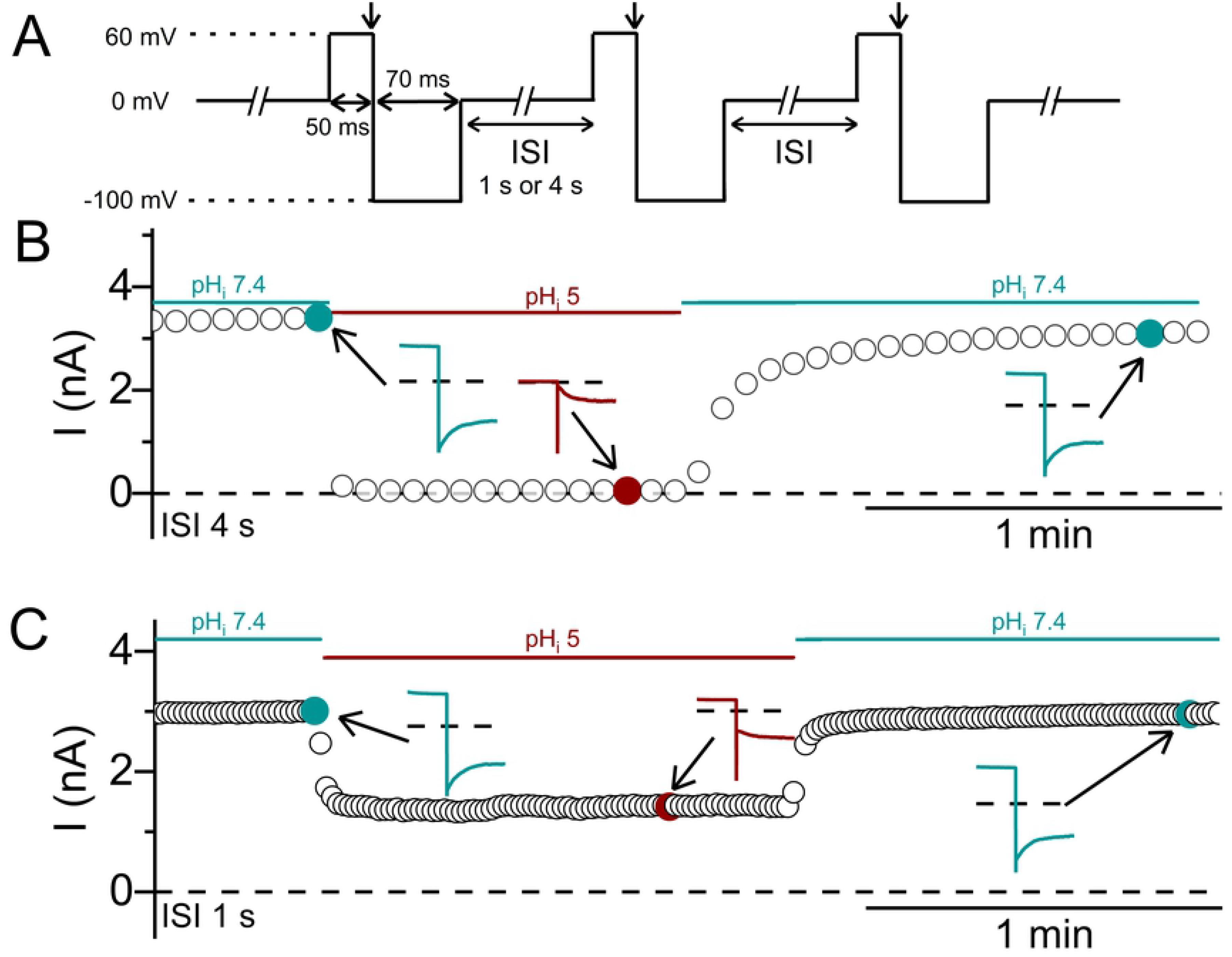
Inhibition of WT CLC-0 by intracellular H^+^. **(A)** Voltage protocol (protocol II) used for the experiments. **(B & C)** Inhibition of the CLC-0 current by an intracellular acidic solution (pHi = 5). Circles represent the current measured at the end of the +60 mV voltage step (downward arrows shown in A). ISI = 4 s and 1 s for the experiments in B and C, respectively. Notice the incomplete inhibition when ISI = 1 sec. Insets show recording traces at indicated time points.

The recordings in Figs. 3 B & C also show that the low pH_i_-induced current inhibition of WT CLC-0 appears to follow an exponentially decaying process. To empirically evaluate the kinetics of the current inhibition by intracellular H^+^, we fit the processes of the low pH_i_-induced current inhibition and the current recovery upon removing low pH_i_ with single-exponential function (Fig. 4 A). The time constants of the current inhibition (τ_inh_) and those of the current recovery (τ_rec_) are plotted against the values of pH_i_ (and also [H^+^]_i_) used for inhibiting the current. Fig. 4 B shows that τ_inh_ is pH_i_-dependent: the higher the [H^+^]_i_, the smaller the value of τ_inh_ (namely, the faster the inhibition). The values of τ_rec_ remain the same in all experiments in Fig. 4 A because they reflect the current recovery to the same final pH_i_ (namely, 7.4).

**FIGURE 4:**
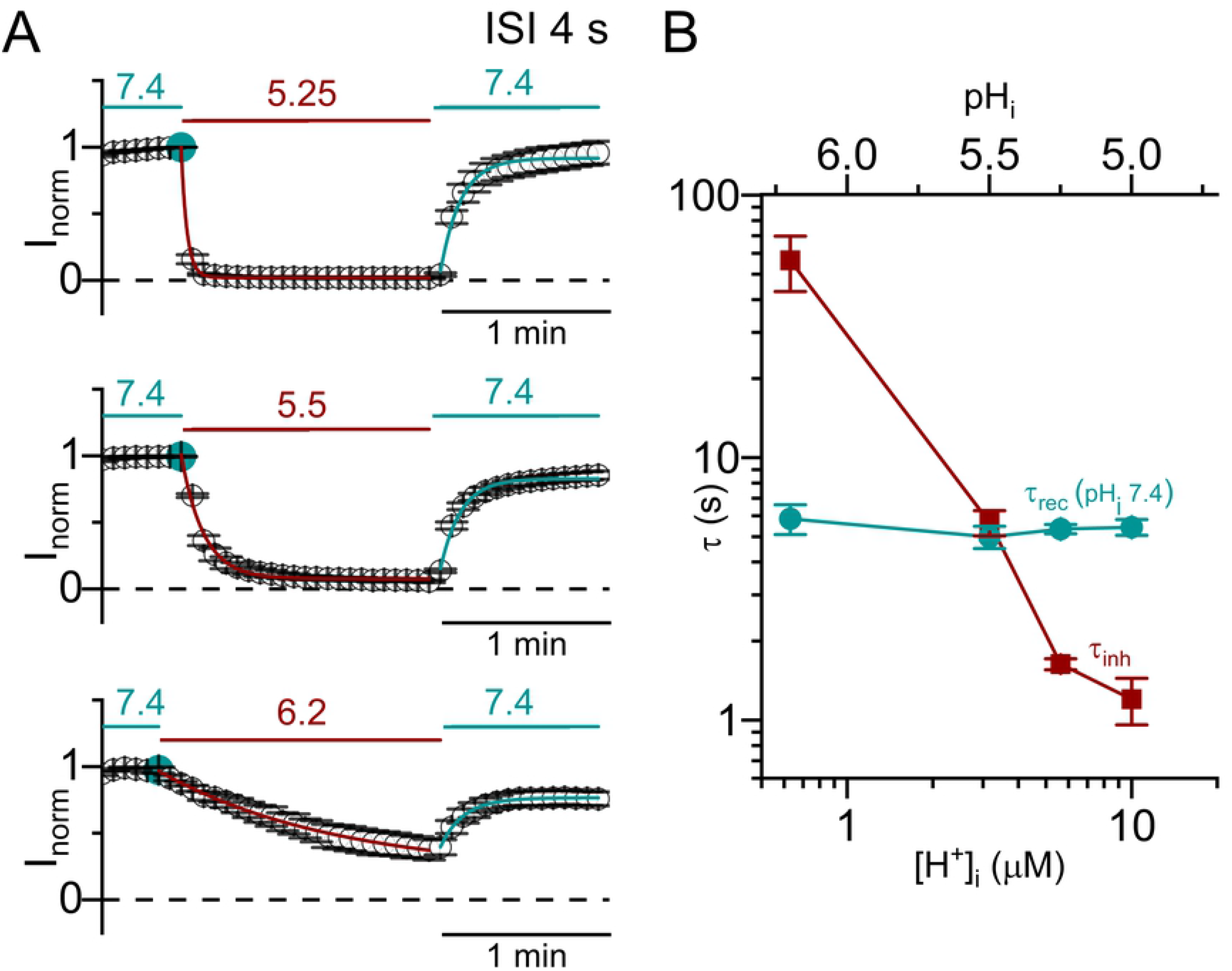
Kinetics of the current inhibition and recovery of WT CLC-0 upon switching pH_i_. Voltage protocol was as that shown in Fig. 3 A. All currents measured at +60 mV were normalized to that obtained right before the application of low pH_i_ solutions. **(A)** Inhibition of WT CLC-0 currents by various low pH_i_ solutions. ISI = 4 s. The numbers above the horizontal lines (teal and red colors) indicate the values of pH_i_. **(B)** Time constants of the inhibition (red squares) plotted against the values of pH_i_ (and thus [H^+^]_i_). Results were obtained from recordings like those shown in A. Time constants of current recovery (at pH_i_ = 7.4) are also plotted (sea green circles).

The experiments in Figs. 3 and 4 were conducted using protocol II with a negative tail voltage step (at −100 mV) which activates the current of WT CLC-0 at low pH_i_. Assessing the effects of membrane voltages on the kinetics of current inhibition and recovery was therefore not accurate. We thus employed a different experimental protocol, namely, altering pH_i_ at constant voltages (protocol III). In such experiments (Fig. 5 A), the current inhibition can still be reasonably fit to a single-exponential function. Nonetheless, no current recovery was observed after switching back to the solution with pH_i_ = 7.4 when the experiment was performed at positive voltages. In comparison with the results obtained with protocol II (Fig. 4), the experiments using protocol III generate a faster current inhibition and a slower current recovery (namely, τ_inh_, is smaller while τ_rec_, is larger). In addition, like those experiments in Fig. 4, τ_inh_ from using protocol III also strongly depends on [H^+^]_i_ (Fig. 5 B) but not on membrane voltages (Fig. 5 C). On the other hand, membrane voltages affect the current recovery significantly in that the more hyperpolarized the membrane voltage, the smaller the value of τ_rec_ (namely, the faster the current recovery rate) (Fig. 5 D).

**FIGURE 5:**
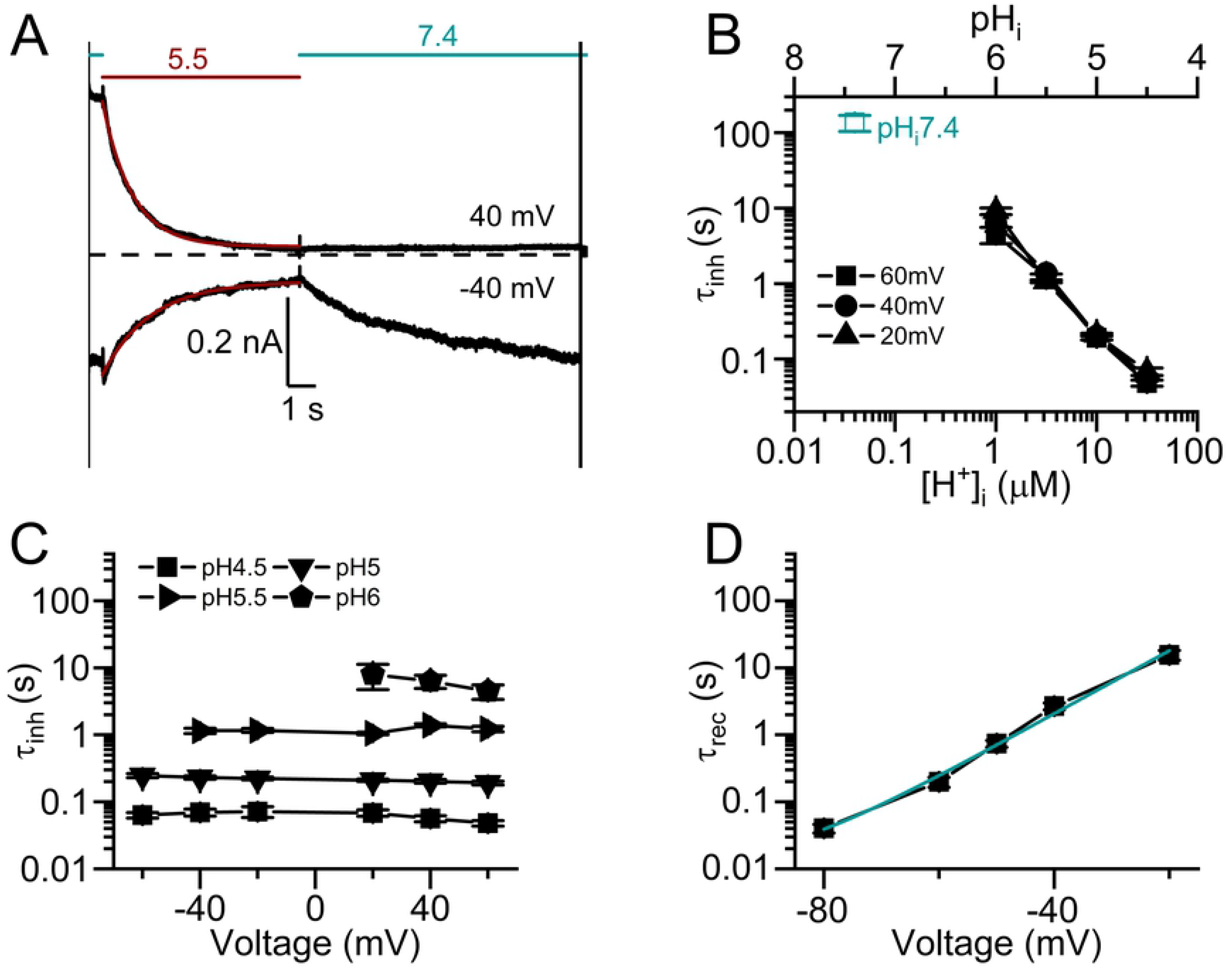
Kinetic analyses of current inhibition and recovery of the H^+^-induced WT CLC-0 inhibition. Experiments were performed with protocol III on excised inside-out patches. Voltages were held constant throughout the recording sweep during which the intracellular solutions with different pH were switched. **(A)** Current inhibition and recovery at ±40 mV. The numbers above the dashed horizontal lines (red and sea green colors) indicate the values of pH_i_. Fitted exponential decay curves (red) are superimposed with the recording traces in black. **(B)** Time constants of inhibition at three voltages (τ_inh_) plotted against [H^+^]_i_ (or pH_i_). The time constant of the slow-gate closure at pH_i_ = 7.4 at room temperature (measured separately) is shown by open square in sea green color. **(C)** Voltage dependence of the inhibition time constant (τ_inh_). **(D)** Current recovery time constants (τ_rec_) against membrane voltages.

It is interesting to note that extrapolating the value of τ_inh_ to a neutral pH_i_ gives a τ_inh_ value of hundreds of sec (Fig. 5 B), which is similar to the relaxation time constant of the CLC-0 slow-gate closure at pH_i_ = 7.4 [38, 39]. This observation suggests an intimate relation between the inhibition of CLC-0 by intracellular H^+^ and the closure of CLC-0’s slow gate. To test this possibility, we examine the intracellular H^+^ inhibition on C212S, a point mutant of CLC-0 in which the slow gate appears to be mostly open [44]. Fig. 6 A & B illustrate the inhibition of WT CLC-0 and C212S by pH_i_ of 5.5 and 4.5, respectively, while the steady-state dose-dependent H^+^ inhibitions between WT CLC-0 and C212S are compared in Fig. 6 C. From this graph, it can be clearly observed that intracellular H^+^ exerts a much weaker inhibitory effect on the C212S mutant than on WT CLC-0.

**FIGURE 6.**
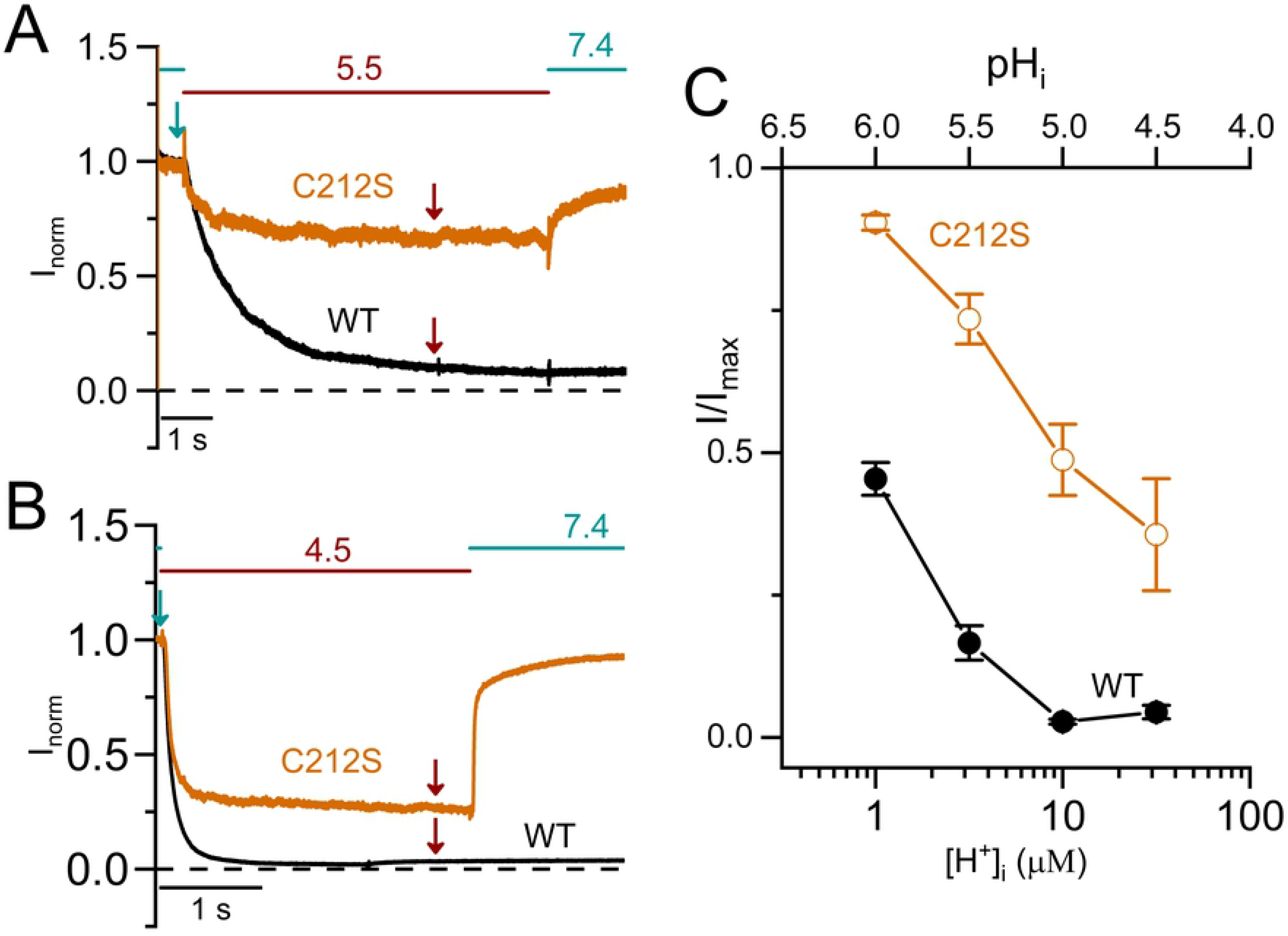
Comparison of low pH_i_-induced inhibitions between WT CLC-0 and the C212S mutant. **(A)** Current inhibition of WT CLC-0 and C212S mutant at +20 mV by pH_i_ = 5.5. **(B)** Current inhibition of WT CLC-0 and C212S mutant at +20 mV by pH_i_ = 4.5. **(C)** Remaining current fraction ( I/I_max_) of WT CLC-0 and C212S mutant against [H^+^]_i_ (or pH_i_). The steady-state current (I) was measured, respectively, at 5 s and 3 s (indicated by wine-colored arrows in A & B, respectively) after applying the pH_i_ 5.5 and pH_i_ 4.5 solutions. I_max_ was the current measured immediately before the low-pH_i_ solution was applied (teal color arrows in A & B).

It has also been well documented that the temperature dependence of the slow gating in WT CLC-0 is significant [38, 39], while that in C212S is weak [44]. The recording traces in Fig. 7 A show that the kinetics of the current inhibition by a low pH_i_ (in this case, pH_i_ = 5.5) is sensitive to temperature in WT CLC-0. To test the temperature dependence of low pH_i_-induced inhibitions in the C212S mutant (Fig. 7 B), pH_i_ = 4.5 was used because this mutant channel is much less sensitive to H^+^ inhibition. Visual inspection of the three recording traces indicates that the temperature dependence of the low pH_i_-induced inhibition in C212S is weak. The averaged results shown in Fig. 7 C reveal a large difference of the temperature dependence of τ_inh_ between WT CLC-0 and the C212S mutant. In Fig. 8 A, the process of the current inhibition by pH_i_ = 4.5 (pink area) and the process of current recovery upon removing high [H^+^]_i_ (light blue area) for WT CLC-0 and the C212S mutant are illustrated. The voltage dependence of the averaged values of τ_inh_ between WT CLC-0 and the C212S mutant are compared in Fig. 8 B, while the comparison of those of τ_rec_ are illustrated in Fig. 8 C. In CLC-0, it is τ_rec_ but not τ_inh_ that is voltage dependent—the more negative the membrane voltage, the smaller the value of τ_rec_ (namely, the faster the recovery from the inhibition). In C212S, a [H^+^]_i_ < 1 μM (namely pH_i_ > 6) generates very little inhibition (Fig. 6), so it is technically necessary to employ very low pH_i_ (4-5) to generate inhibition for the experiments. The value of τ_inh_ is small (fast current inhibition) likely because of the high [H^+^]_i_ at pH_i_ = 4.5. Interestingly, τ_rec_ is also small in C212S, reflecting a faster current recovery process than that in WT CLC-0. Fig. 8 B & C show that the voltage dependence of τ_inh_ and τ_rec_ are both weak for the C212S mutant.

**FIGURE 7:**
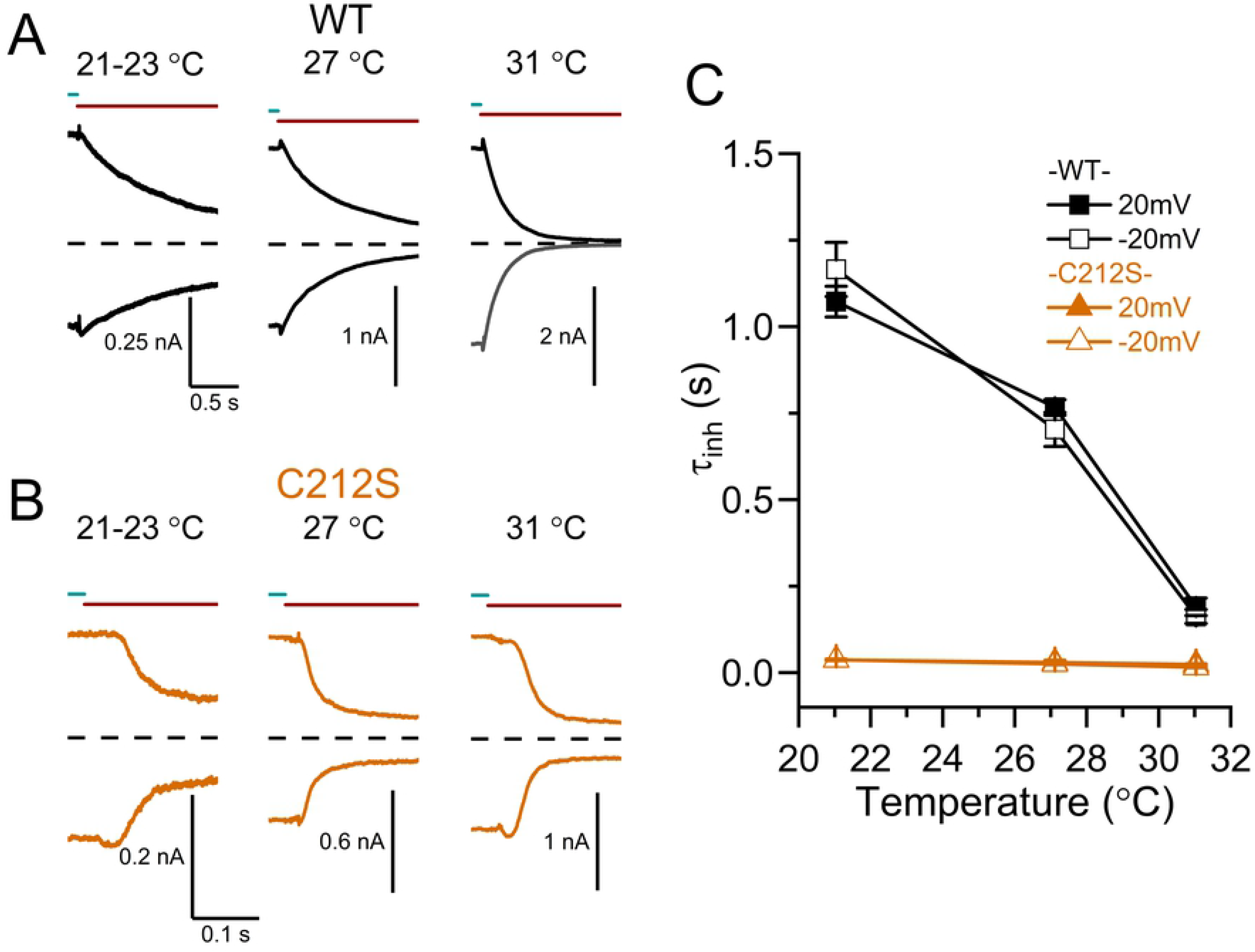
Comparing the temperature dependence of low pH_i_-induced inhibitions between WT CLC-0 and the C212S mutant. **(A)** Inhibitions of WT CLC-0 by a solution with pH_i_ = 5.5 at three temperatures. **(B)** Inhibitions of the C212S mutant by a solution with pH_i_ = 4.5. All recording traces in A & B were obtained at V_m_ = ±20 mV. **(C)** Time constants of H^+^ inhibition of WT CLC-0 and the C212S mutant plotted against temperature.

**FIGURE 8.**
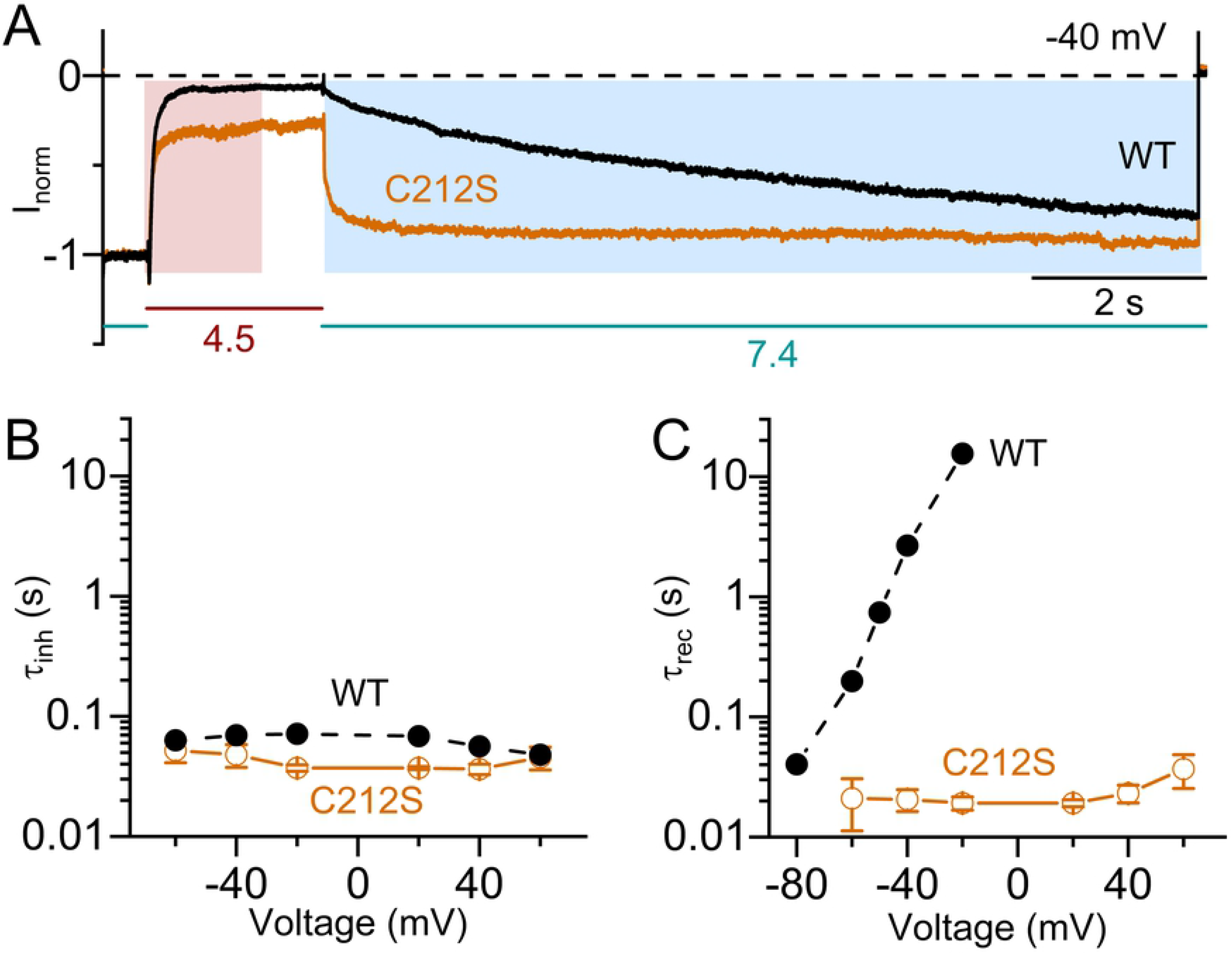
Comparing the voltage dependence of low pH_i_-induced inhibition between WT CLC-0 and the C212S mutant. **(A)** Recording traces showing the current inhibition induced by a solution with pH_i_ = 4.5 in WT CLC-0 and C212S. V_m_ = −40 mV. Values of the pH_i_ were shown below the colored horizontal lines. The values of τ_inh_ and τ_rec_ were obtained by fitting the current inhibition process (region shaded in pink color) and the current recovery process (region shaded in light blue) to single-exponential functions. **(B)** Voltage dependence of τ_inh_ of WT CLC-0 and C212S. All data points were obtained from the inhibition induced by a solution with pH_i_ = 4.5. **(C)** Voltage dependence of τ_rec_ of WT CLC-0 and C212S after pH_i_ = 4.5 was switched back to pH_i_ = 7.4. At positive voltages, current recovery was observed in the C212S mutant but not in WT CLC-0.

If the lower sensitivity to intracellular H^+^ inhibition in C212S is due to a reduced slow-gate closure in this mutant, other mutations that also prevent the channel from closing the slow gate may exhibit similar low sensitivity to intracellular H^+^ inhibition. In Fig. 9 A, we compare the inhibition by low pH_i_ solutions (pH_i_ = 5.5 and 4.5 in the upper and lower panel, respectively) between WT CLC-0 and another mutant, Y512A, which has also been shown to largely prevent the slow gate from closing [45]. The steady-state [H^+^]_i_-dependent current inhibition in Fig. 9 B indeed shows that the mutant Y512A, like C212S, is also more resistant to the intracellular H^+^ inhibition than WT CLC-0.

**FIGURE 9:**
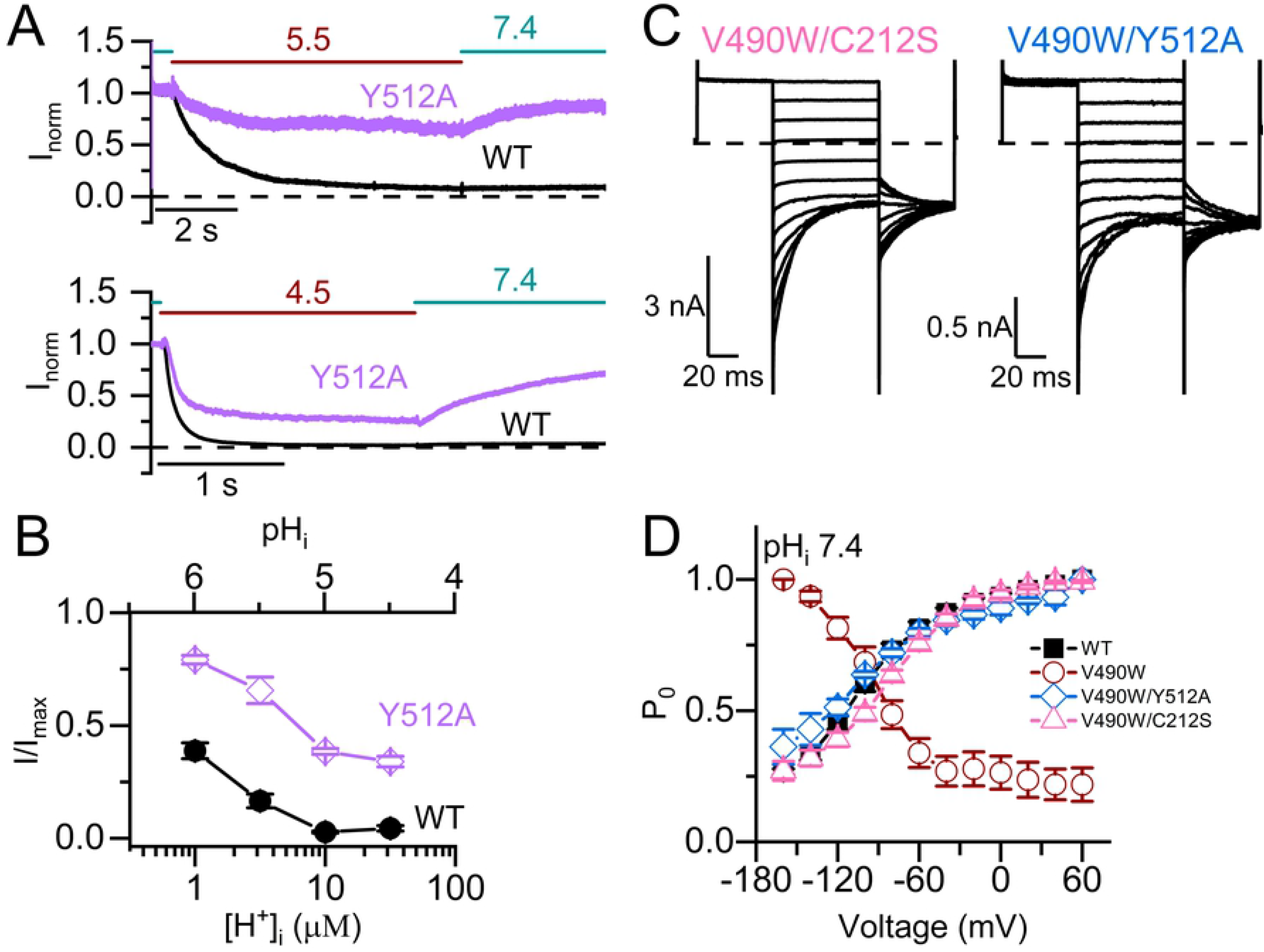
Correcting the inverted voltage-dependent opening in the V490W mutant by mutations that inhibit slow-gate closure. **(A)** Comparing the inhibitions of WT CLC-0 and the Y512A mutant by solutions with pH_i_ = 5.5 (upper panel) and 4.5 (lower panel). **(B)** Remaining current fractions of WT CLC-0 and the Y512A mutant after the current inhibition by various [H^+^]_i_. **(C)** Recording traces of the double mutants V490W/C212S (left) and V490W/Y512A (right) obtained using the experimental protocol I with the tail step voltage at −100 mV. In both recordings, pH_i_ = 7.4. **(D)** Relative P_o_ of WT CLC-0, three mutants, C212S, V490W/C212S and V490W/Y512A. All data were obtained at pH_i_ =7.4.

On the other hand, if the inverted voltage dependent opening of the V490W mutant is due to an excessive slow-gate closure in neutral pH_i_, the C212S mutation or the Y512A mutation may reduce the excessive slow-gate closure caused by the V490W mutation. In Fig. 9 C, we show the recording traces of two double mutants, V490W/C212S and V490/Y512A, at pH_i_ = 7.4, using the voltage protocol I shown in Fig. 2 A. In Fig. 9 D, we plot the normalized instantaneous tailed currents of WT CLC-0 (black solid squares), V490W (red circles), the V490W/C212S mutant (triangles), and the V490W/Y512A mutant (diamonds) obtained from recordings at pH_i_ = 7.4. It can be seen that the inverted voltage-dependent opening in the V490W mutant disappears in both the V490W/C212S and V490W/Y512A double mutants. The voltage dependent opening of these two double mutants looks very similar to that of WT CLC-0. These results suggest that the inverted voltage dependent opening of the V490W mutant may be due to an excessive slow-gate closure at the neutral pH_i_.

## Discussion

The two types of CLC family members, Cl^−^ channels (such as CLC-0, CLC-1, CLC-2 and CLC-Ks) and Cl^−^/H^+^ antiporters (such as bacterial CLCs and mammalian CLCs expressed in intracellular organelles), are structurally similar to each other [17–22], so the gating mechanisms of CLC Cl^−^ channels are thought to be driven by a background Cl^−^/H^+^ antiporter activity [52]. Indeed, a change of pH close to the cell membrane has been demonstrated during the opening and closing of CLC channels [12]. In CLC-0, Cl^−^ and H^+^ both increase P_o_^f^ [27, 28, 35, 36]. Mechanistically, protonation of the E-gate is thought to help swing the fast gate away from the pore while Cl^−^ competes with the E-gate for the Cl^−^ binding site S_ext_. On the other hand, the molecular mechanism of slow gating of CLC-0 remains a mystery. At the single-channel level, Richard and Miller [40] discovered a “non-equilibrium” gating cycle for CLC-0’s slow gating. The phenomenon involves an asymmetry in the transitions between the slow-gate open state and the inactivation state: the channels are more likely to enter the inactivation state from the one-pore open state while leaving the inactivation state to the two-pore open state. They demonstrated that this non-equilibrium gating is facilitated by a transmembrane Cl^−^ flux. Later experiments by Lisal and Maduke [41] discovered that the H^+^ gradient across the membrane may be an even more powerful energy source to promote this non-equilibrium gating. The structural-functional basis underlying these findings of non-equilibrium gating remains unsolved.

The results from our present work reveal a potent inhibitory effect on CLC-0 currents by intracellular H^+^. An intracellular solution with pH_i_ = 6 ([H^+^]_i_ = 1 μM) inhibits the steady-state current of the WT channel by ~50 % (Fig. 6 C). However, the kinetics of the H^+^-induced inhibition is slow. Hence, we suspected that the inhibition of CLC-0 by intracellular H^+^ is related to the slow-gate closure. The dependence of this H^+^ inhibition on membrane voltage and temperature is also consistent with the properties of CLC-0’s slow gating—the more negative the membrane voltage, the faster the current recovery from the inhibition [53], and the higher the temperature, the faster the inhibition relaxation kinetics [38, 39]. Furthermore, inhibition effects by intracellular H^+^ on the C212S and Y512A mutants, in which the slow gates are mostly open [44, 45], were much weaker than that on the WT CLC-0. These findings indicate that intracellular H^+^ enhances the closure of the slow gate of CLC-0, thus generating the current inhibition.

So far, most of the proposed slow-gating mechanisms of CLC-0 are vague. For example, a conformational change of the channel has been suggested to be involved in the slow-gating mechanism of CLC-0 [43, 46]. Yet, it is not known what exact conformational change is and whether the speculated conformational change is causally involved in the slow gating. A more specific slow-gate closing mechanism was recently proposed by Bennetts and Parker [45]. Based on the observation that the slow-gate closure appears not present in the Y512A mutant of CLC-0, they proposed a “pincer” occlusion near S_cen_ as the slow-gate closing mechanism, where the carboxylate of the E-gate (E166 in CLC-0) forms a hydrogen bond with the phenolic hydroxyl group of a tyrosine residue (Y512 in CLC-0). However, as Jentsch and Pusch point out in their recent review [1], such interaction is not possible because manipulation of the corresponding tyrosine residue in another CLC channel homologue, CLC-K (Y520A mutation in CLC-K), still produces a similar gating change [54]. While lacking the gating glutamate, CLC-K instead has a valine residue at the equivalent position of E166 of CLC-0. Furthermore, the proposal of Bennetts and Parker would have predicted that a lower pH_i_, which favors protonation of the sidechain of the E-gate, would interfere with hydrogen bond formation between E166 and Y512, a scenario directly opposite to that observed in this study.

Although our study does not illustrate the structural mechanism of the slow gating, the intracellular H^+^ inhibition of CLC-0 does provide several insights for this gating mechanism. First, we found a strong dependence of the inhibition rate on [H^+^]_i_. At a room temperature of 21-22 ̊C, the values of τ_inh_ from experiments using protocol II are ~ 0.1, 1 and ~10 sec at pH_i_ = 4.5, 5.5, and 6, respectively (Fig. 5 B). This finding explains the previous observation of the very large (several hundred seconds) relaxation time constant of the slow-gate closure at a neutral pH_i_ of 7.4 ([H^+^]_i_ < 100 nM) at the same temperature [38, 39]. Thus, the binding of intracellular H^+^ to CLC-0 likely initiates the process of slow-gate closure in CLC-0.

The second insight our study offers is to explain the inverted voltage-dependent activation in some CLC channel mutants (such as the V490W mutant shown in this study). Unlike WT CLC-0, membrane depolarization is unable to open this mutant. On the other hand, this mutant is opened by membrane hyperpolarization (Fig. 9 D), a voltage dependence similar to that of the slow-gate opening of CLC-0. We also show that intracellular H^+^ speeds up the rate of the slow-gate closure, thus generating an apparent inverted voltage-dependent channel opening even in WT CLC-0. It is thus possible that mutants of CLC-0 with inverted voltage-dependent activation have a fast slow-gate closure at neutral pH_i_. By constructing the V490W mutation in the background of C212S or Y512A, two mutants with little slow-gate closure, we show the double mutants, V490W/C212S and V490W/Y512A, no longer exhibit the inverted voltage-dependent opening (Fig. 9 C & D).

Careful inspection of the intracellular H^+^ inhibition of the C212S and Y512A mutants may provide further insight into the mechanism of the slow gating. The slow gate in these two mutants are mostly open. Accordingly, these two mutations significantly weaken the effects of manipulations that facilitate the slow-gate closure in CLC-0, such as a rise of experimental temperature or an application of extracellular zinc ions (Zn^2+^) [44, 45]. Here we add another manipulation that facilitates the inactivation in WT CLC-0, namely, an increase of [H^+^]_i_. However, both C212S and Y512A are still inhibited by intracellular H^+^, although a much higher concentration of [H^+^]_i_ is required. The rates of current inhibition in these two mutants upon applying [H^+^]_i_ seem to be roughly similar to that of WT CLC-0 (Fig. 8 A & B). On the other hand, current recovery from H^+^ inhibition is much faster in these two mutants than in WT CLC-0. Thus, a weaker H^+^ inhibition in these mutants is due to a faster current recovery (Fig. 8 A & C). We thus propose the following model (model 1) to explain the H^+^-induced slow-gate closure of CLC-0:

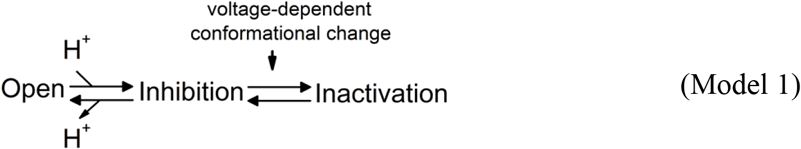

Here we propose that after the binding of intracellular H^+^ to the channel, the channel pore is closed (“Inhibition” state), followed by the channel’s entry into a stable “Inactivation” state via a voltage-dependent gating step. The H^+^ titration curves show that the apparent pK_a_ (namely, the pH_i_ where the current is half inhibited by H^+^) of WT CLC-0 is ~ 6, while the apparent pK_a_’s of C212S and Y512A are about 1 pH unit lower (Fig. 6 C & Fig. 9 B). Since the pK_a_ changes by C212S and Y512A mutations are similar, it is less likely that the mutation effects are caused by a lack of H^+^ titration of the thiol group of C212S and the phenolic hydroxyl group of Y512 (which have very different true pK_a_ values). We suggest that the C212S and Y512A mutations do not alter the H^+^ binding step in model 1 but significantly increase the energy barrier between the “Inhibition” state and the “Inactivation” state, thus preventing the channel from entering the “Inactivation” state. Because these two mutant channels are rarely in the stable “Inactivation” state even with high [H^+^]_i_, the current recovery upon removing H^+^ is fast and not voltage dependent.

Interesting questions remain regarding how intracellular H^+^ binding to the channel from the intracellular side inhibits the Cl^−^ current flow in CLC-0, and how C212S and Y512A mutations prevent the channel from entering the inactivation state. Answering these questions are not possible without further specific experiments. For example, we have not examined the functional roles of Cl^−^ in this H^+^ inhibitory effect on CLC-0’s slow gating. The present study also does not address the question whether transmembrane [H^+^] gradient plays a role in the low-pH_i_-induced inhibition. Given the significant difference in the current recovery between the positive and negative voltages (Fig. 5 A), it is interesting to understand whether this difference results from a true voltage-dependent effect or is related to the direction of Cl^−^ or H^+^ flux. Furthermore, mutations of amino acid residues with a titratable sidechain will help us identify the potential H^+^-binding site(s) responsible for the intracellular H^+^-induced current inhibition, and this endeavor may help us explore what specific conformational change is involved in slow gating. Previous research suggested that the mutations, at least in the case of C212S, may render the relative movement of the two subunits less likely [46]. This scenario is certainly consistent with the model proposed above if such a conformational change is associated with the gating step between the “Inhibition” and the “Inactivation” state.

In summary, although the molecular mechanism underlying the slow gating of CLC-0 is still largely unknown, the experimental results we present in this study offer several refinements of our knowledge. First, we find that increased [H^+^]_i_ increases the rate of slow-gate closure of WT CLC-0, and thus the slow kinetics (tens to hundreds of seconds) of CLC-0 slow gating at neutral pH_i_ is likely due to low [H^+^]_i_ (< 0.1 μM) rather than due to a large conformational change of the channel protein. Second, the inverted voltage-dependent activation of the V490W mutant is likely due to an excessively fast slow-gate closure of the mutant at neutral pH_i_. Finally, our results are still consistent with an involvement of a protein conformational change in the slow gating, which likely occurs between the “Inhibition” and the “Inactivation” states shown in model 1. Future experiments based on these insights may help us further unveil the slow-gating mechanism of CLC channels.

## Acknowledgment

We thank Dr. Wei-Ping Yu for constructing the cDNA constructs of the WT CLC-0 and all the mutants.

## Author Contributions

Conceived and designed the experiments: HCK, YY, TC. Performed the electrophysiological experiments: HCK, YY. Analyzed the data, HCK, YY; Contributed reagents/materials/analysis tools: TC. Wrote the paper: HCK, RHF, TC.

## References

1. Jentsch TJ, Pusch M. CLC Chloride Channels and Transporters: Structure, Function, Physiology, and Disease. Physiol Rev. 2018;98(3):1493–590 doi: 10.1152/physrev.00047.2017. PubMed PMID: 29845874.

2. Jentsch TJ. Discovery of CLC transport proteins: cloning, structure, function and pathophysiology. J Physiol. 2015;593(18):4091–109 doi: 10.1113/JP270043. PubMed PMID: 25590607; PubMed Central PMCID: PMCPMC4594286.

3. Jentsch TJ, Poet M, Fuhrmann JC, Zdebik AA. Physiological functions of CLC Cl-channels gleaned from human genetic disease and mouse models. Annu Rev Physiol. 2005;67:779–807. PubMed PMID: 15709978.

4. Jentsch TJ, Steinmeyer K, Schwarz G. Primary structure of Torpedo marmorata chloride channel isolated by expression cloning in Xenopus oocytes. Nature. 1990;348(6301):510–4 PubMed PMID: 2174129.

5. Steinmeyer K, Ortland C, Jentsch TJ. Primary structure and functional expression of a developmentally regulated skeletal muscle chloride channel. Nature. 1991;354(6351):301–4 PubMed PMID: 1659664.

6. Thiemann A, Grunder S, Pusch M, Jentsch TJ. A chloride channel widely expressed in epithelial and non-epithelial cells. Nature. 1992;356(6364):57–60 PubMed PMID: 1311421.

7. Uchida S, Sasaki S, Furukawa T, Hiraoka M, Imai T, Hirata Y, et al. Molecular cloning of a chloride channel that is regulated by dehydration and expressed predominantly in kidney medulla. J Biol Chem. 1993;268(6):3821–4 PubMed PMID: 7680033.

8. Maduke M, Pheasant DJ, Miller C. High-level expression, functional reconstitution, and quaternary structure of a prokaryotic ClC-type chloride channel. J Gen Physiol. 1999;114(5):713–22 PubMed PMID: 10539975.

9. Accardi A, Miller C. Secondary active transport mediated by a prokaryotic homologue of ClC Cl-channels. Nature. 2004;427(6977):803–7 PubMed PMID: 14985752.

10. Iyer R, Iverson TM, Accardi A, Miller C. A biological role for prokaryotic ClC chloride channels. Nature. 2002;419(6908):715–8 PubMed PMID: 12384697.

11. Lloyd SE, Gunther W, Pearce SH, Thomson A, Bianchi ML, Bosio M, et al. Characterisation of renal chloride channel, CLCN5, mutations in hypercalciuric nephrolithiasis (kidney stones) disorders. Hum Mol Genet. 1997;6(8):1233–9 PubMed PMID: 9259268.

12. Picollo A, Pusch M. Chloride/proton antiporter activity of mammalian CLC proteins ClC-4 and ClC-5. Nature. 2005;436(7049):420–3 PubMed PMID: 16034421.

13. Poet M, Kornak U, Schweizer M, Zdebik AA, Scheel O, Hoelter S, et al. Lysosomal storage disease upon disruption of the neuronal chloride transport protein ClC-6. Proc Natl Acad Sci U S A. 2006;103(37):13854–9 PubMed PMID: 16950870.

14. Matsumoto A, Matsui I, Mori T, Sakaguchi Y, Mizui M, Ueda Y, et al. Severe Osteomalacia with Dent Disease Caused by a Novel Intronic Mutation of the CLCN5 gene. Intern Med. 2018;57(24):3603–10 doi: 10.2169/internalmedicine.1272-18. PubMed PMID: 30101934; PubMed Central PMCID: PMCPMC6355425.

15. Piwon N, Gunther W, Schwake M, Bosl MR, Jentsch TJ. ClC-5 Cl--channel disruption impairs endocytosis in a mouse model for Dent’s disease. Nature. 2000;408(6810):369–73 PubMed PMID: 11099045.

16. Middleton RE, Pheasant DJ, Miller C. Purification, reconstitution, and subunit composition of a voltage-gated chloride channel from Torpedo electroplax. Biochemistry. 1994;33(45):13189–98 PubMed PMID: 7947726.

17. Dutzler R, Campbell EB, Cadene M, Chait BT, MacKinnon R. X-ray structure of a ClC chloride channel at 3.0 A reveals the molecular basis of anion selectivity. Nature. 2002;415(6869):287–94 PubMed PMID: 11796999.

18. Dutzler R, Campbell EB, MacKinnon R. Gating the selectivity filter in ClC chloride channels. Science. 2003;300(5616):108–12 PubMed PMID: 12649487.

19. Feng L, Campbell EB, Hsiung Y, MacKinnon R. Structure of a eukaryotic CLC transporter defines an intermediate state in the transport cycle. Science. 2010;330(6004):635–41 Epub 2010/10/12. doi: science.1195230 [pii]10.1126/science.1195230. PubMed PMID: 20929736; PubMed Central PMCID: PMC3079386.

20. Park E, Campbell EB, MacKinnon R. Structure of a CLC chloride ion channel by cryo-electron microscopy. Nature. 2017;541(7638):500–5 doi: 10.1038/nature20812. PubMed PMID: 28002411; PubMed Central PMCID: PMCPMC5576512.

21. Park E, MacKinnon R. Structure of the CLC-1 chloride channel from Homo sapiens. Elife. 2018;7. doi: 10.7554/eLife.36629. PubMed PMID: 29809153; PubMed Central PMCID: PMCPMC6019066.

22. Wang K, Preisler SS, Zhang L, Cui Y, Missel JW, Gronberg C, et al. Structure of the human ClC-1 chloride channel. PLoS Biol. 2019;17(4):e3000218. doi: 10.1371/journal.pbio.3000218. PubMed PMID: 31022181; PubMed Central PMCID: PMCPMC6483157.

23. Middleton RE, Pheasant DJ, Miller C. Homodimeric architecture of a ClC-type chloride ion channel. Nature. 1996;383(6598):337–40 PubMed PMID: 8848046.

24. Miller C. Open-state substructure of single chloride channels from Torpedo electroplax. Philos Trans R Soc Lond B Biol Sci. 1982;299(1097):401–11 PubMed PMID: 6130538.

25. Miller C, White MM. Dimeric structure of single chloride channels from Torpedo electroplax. Proc Natl Acad Sci U S A. 1984;81(9):2772–5 PubMed PMID: 6326143.

26. Chen TY. Structure and function of clc channels. Annu Rev Physiol. 2005;67:809–39. Epub 2005/02/16. doi: 10.1146/annurev.physiol.67.032003.153012. PubMed PMID: 15709979.

27. Pusch M, Ludewig U, Rehfeldt A, Jentsch TJ. Gating of the voltage-dependent chloride channel CIC-0 by the permeant anion. Nature. 1995;373(6514):527–31 PubMed PMID: 7845466.

28. Chen TY, Miller C. Nonequilibrium gating and voltage dependence of the ClC-0 Cl-channel. J Gen Physiol. 1996;108(4):237–50 PubMed PMID: 8894974.

29. Chen TY, Chen MF, Lin CW. Electrostatic control and chloride regulation of the fast gating of ClC-0 chloride channels. J Gen Physiol. 2003;122(5):641–51 Epub 2003/10/29. doi: 10.1085/jgp.200308846. PubMed PMID: 14581587; PubMed Central PMCID: PMCPMC2229583.

30. Chen TY. Coupling gating with ion permeation in ClC channels. Sci STKE. 2003;2003(188):pe23. Epub 2003/06/26. doi: 10.1126/stke.2003.188.pe23. PubMed PMID: 12824475.

31. Dutzler R. Structural basis for ion conduction and gating in ClC chloride channels. FEBS Lett. 2004;564(3):229–33 PubMed PMID: 15111101.

32. Engh AM, Faraldo-Gomez JD, Maduke M. The mechanism of fast-gate opening in ClC-0. J Gen Physiol. 2007;130(4):335–49 Epub 2007/09/12. doi: jgp.200709759 [pii]10.1085/jgp.200709759. PubMed PMID: 17846164; PubMed Central PMCID: PMC2151655.

33. Zifarelli G, Murgia AR, Soliani P, Pusch M. Intracellular proton regulation of ClC-0. J Gen Physiol. 2008;132(1):185–98 PubMed PMID: 18591423.

34. Zifarelli G, Pusch M. The role of protons in fast and slow gating of the Torpedo chloride channel ClC-0. Eur Biophys J. 2010;39(6):869–75 doi: 10.1007/s00249-008-0393-x. PubMed PMID: 19132363.

35. Hanke W, Miller C. Single chloride channels from Torpedo electroplax. Activation by protons. J Gen Physiol. 1983;82(1):25–45 PubMed PMID: 6310023.

36. Chen MF, Chen TY. Different fast-gate regulation by external Cl(−) and H(+) of the muscle-type ClC chloride channels. J Gen Physiol. 2001;118(1):23–32 PubMed PMID: 11429442.

37. White MM, Miller C. A voltage-gated anion channel from the electric organ of Torpedo californica. J Biol Chem. 1979;254(20):10161–6 PubMed PMID: 489590.

38. Pusch M, Ludewig U, Jentsch TJ. Temperature dependence of fast and slow gating relaxations of ClC-0 chloride channels. J Gen Physiol. 1997;109(1):105–16 PubMed PMID: 8997669.

39. Chen TY. Extracellular Zinc Ion Inhibits ClC-0 Chloride Channels by Facilitating Slow Gating. J Gen Physiol. 1998;112(6):715–26 PubMed PMID: 9834141.

40. Richard EA, Miller C. Steady-state coupling of ion-channel conformations to a transmembrane ion gradient. Science. 1990;247(4947):1208–10 PubMed PMID: 2156338.

41. Lisal J, Maduke M. The ClC-0 chloride channel is a ‘broken’ Cl-/H+ antiporter. Nat Struct Mol Biol. 2008;15(8):805–10 Epub 2008/07/22. doi: nsmb.1466 [pii]10.1038/nsmb.1466. PubMed PMID: 18641661; PubMed Central PMCID: PMC2559860.

42. Fong P, Rehfeldt A, Jentsch TJ. Determinants of slow gating in ClC-0, the voltage-gated chloride channel of Torpedo marmorata. Am J Physiol. 1998;274(4 Pt 1):C966–73. PubMed PMID: 9575793.

43. Yu Y, Tsai MF, Yu WP, Chen TY. Modulation of the slow/common gating of CLC channels by intracellular cadmium. J Gen Physiol. 2015;146(6):495–508 doi: 10.1085/jgp.201511413. PubMed PMID: 26621774; PubMed Central PMCID: PMCPMC4664824.

44. Lin YW, Lin CW, Chen TY. Elimination of the slow gating of ClC-0 chloride channel by a point mutation. J Gen Physiol. 1999;114(1):1–12 PubMed PMID: 10398688.

45. Bennetts B, Parker MW. Molecular determinants of common gating of a ClC chloride channel. Nat Commun. 2013;4:2507. Epub 2013/09/26. doi: ncomms3507 [pii]10.1038/ncomms3507. PubMed PMID: 24064982.

46. Bykova EA, Zhang XD, Chen TY, Zheng J. Large movement in the C terminus of CLC-0 chloride channel during slow gating. Nat Struct Mol Biol. 2006;13(12):1115–9 Epub 2006/11/23. doi: 10.1038/nsmb1176. PubMed PMID: 17115052.

47. Bretag AH. Muscle chloride channels. Physiol Rev. 1987;67(2):618–724 Epub 1987/04/01. PubMed PMID: 2436244.

48. Chen TY. Single myotonia mutation strikes multiple mechanisms of a chloride channel. J Physiol. 2012;590(Pt 15):3407. Epub 2012/08/03. doi: 10.1113/jphysiol.2012.238337. PubMed PMID: 22855049; PubMed Central PMCID: PMCPMC3547255.

49. Fahlke C, Rudel R, Mitrovic N, Zhou M, George AL, Jr. An aspartic acid residue important for voltage-dependent gating of human muscle chloride channels. Neuron. 1995;15(2):463–72 PubMed PMID: 7646898.

50. Macias MJ, Teijido O, Zifarelli G, Martin P, Ramirez-Espain X, Zorzano A, et al. Myotonia-related mutations in the distal C-terminus of ClC-1 and ClC-0 chloride channels affect the structure of a poly-proline helix. Biochem J. 2007;403(1):79–87 Epub 2006/11/17. doi: BJ20061230 [pii]10.1042/BJ20061230. PubMed PMID: 17107341; PubMed Central PMCID: PMC1828897.

51. Lin CW, Chen TY. Cysteine modification of a putative pore residue in ClC-0: implication for the pore stoichiometry of ClC chloride channels. J Gen Physiol. 2000;116(4):535–46 PubMed PMID: 11004203.

52. Miller C. ClC chloride channels viewed through a transporter lens. Nature. 2006;440(7083):484–9 PubMed PMID: 16554809.

53. Miller C, White MM. A voltage-dependent chloride conductance channel from Torpedo electroplax membrane. Ann N Y Acad Sci. 1980;341:534–51. PubMed PMID: 6249158.

54. Gradogna A, Pusch M. Alkaline pH block of CLC-K kidney chloride channels mediated by a pore lysine residue. Biophys J. 2013;105(1):80–90 doi: 10.1016/j.bpj.2013.05.044. PubMed PMID: 23823226; PubMed Central PMCID: PMCPMC3699751.

